# Analytical Validation of Multiplex Biomarker Assay to Stratify Colorectal Cancer Samples into Molecular Subtypes

**DOI:** 10.1101/174847

**Authors:** Chanthirika Ragulan, Katherine Eason, Elisa Fontana, Gift Nyamundanda, Yatish Patil, Pawan Poudel, Rita T. Lawlor, Maguy Del Rio, Koo Si-Lin, Tan Wah Siew, Francesco Sclafani, Ruwaida Begum, Larissa S. Teixeira Mendes, Pierre Martineau, Aldo Scarpa, Iain Beehuat Tan, David Cunningham, Anguraj Sadanandam

## Abstract

Previously, we classified colorectal cancers (CRCs) into five CRCA subtypes with different prognoses and potential treatment responses, using a 786-gene signature. We merged our subtypes and those described by five other groups into four consensus molecular subtypes (CMS) that are similar to CRCA subtypes. Here we demonstrate the analytical development and application of a custom NanoString platform-based biomarker assay to stratify CRC into subtypes. To reduce costs, we switched from the standard protocol to a custom modified protocol (NanoCRCA) with a high Pearson correlation coefficient (>0.88) between protocols. Technical replicates were highly correlated (>0.96). The assay included a reduced robust 38-gene panel from the 786-gene signature that was selected using an in-laboratory developed computational pipeline of class prediction methods. We applied our NanoCRCA assay to untreated CRCs including fresh-frozen and formalin-fixed paraffin-embedded (FFPE) samples (n=81) with matched microarray or RNA-Seq profiles. We further compared the assay results with CMS classification, different platforms (microarrays/RNA-Seq) and gene-set classifiers (38 and 786 genes). NanoCRCA classified fresh-frozen samples (n=39; not including those showing a mixture of subtypes) into all five CRCA subtypes with overall high concordance across platforms (89.7%) and with CMS subtypes (84.6%), independent of tumour cellularity. This analytical validation of the assay shows the association of subtypes with their known molecular, mutational and clinical characteristics. Overall, our modified NanoCRCA assay with further clinical assessment may facilitate prospective validation of CRC subtypes in clinical trials and beyond.

**Novelty and Impact:** We previously identified five gene expression-based CRC subtypes with prognostic and potential predictive differences using a 786-gene signature and microarray platform. Subtype-driven clinical trials require a validated assay suitable for routine clinical use. This study demonstrates, for the first time, how molecular CRCA subtype can be detected using NanoString Technology-based biomarker assay (NanoCRCA) suitable for clinical validation. NanoCRCA is suitable for analysing FFPE samples, and this assay may facilitate patient stratification within clinical trials.

## Introduction

Colorectal cancer (CRC) is the fourth leading cause of cancer-related deaths worldwide ^1^. The median overall survival of metastatic (m)CRC patients with unresectable disease remains around 24 months with standard chemotherapies. Targeted therapies including anti-EGFR antibodies, anti-angiogenic and immunotherapy agents may extend survival up to 30 months in selected patients ^2^. However, how to identify patients who will benefit from different drug options remains challenging. Additional predictive biomarkers are required to spare patients from unnecessary toxicities, improve outcomes and increase cost-effectiveness of treatment.

In order to classify colorectal cancers into subtypes with distinct biology, helping to effectively match existing therapies and facilitate subtype-specific therapeutic development, we previously identified five distinctive gene expression subtypes using a 786-gene signature^3^. Based on the gene expression similarities with different cell types of the normal colonic mucosa, the 5 subtypes were named goblet-like, enterocyte, stem-like, inflammatory and transit-amplifying (TA). We demonstrated significantly poorer disease-free survival (DFS) in untreated patients for the stem-like subtype, intermediate DFS for inflammatory and enterocyte, and better DFS for goblet-like and TA. Then, from two different datasets that included drug response information, we observed increased responses within the stem-like subtype to irinotecan, fluorouracil and leucovorin treatment combination (FOLFIRI) and the TA subtypes to anti-EGFR monoclonal antibody (cetuximab) ^3-5^. These treatment responses were further validated by other studies ^6, 7^.

Five other groups independently identified between 3 and 6 molecularly distinct CRC subtypes based on expression profiles ^8-12^. These and our findings were aggregated by a CRC Subtyping Consortium (CRCSC) into 4 consensus molecular subtypes (CMS): CMS1 (similar to inflammatory subtype); CMS2 (enterocyte and TA); CMS3 (goblet-like); and CMS4 (stem-like), plus a “mixed” subtype representing either the existence of additional subtypes or the presence of multiple subtypes in a single sample ^13^. In the current manuscript, we found “mixed” subtypes to have multiple existing subtypes. CRCA and CMS subtypes are similar except that the enterocyte and TA subtypes of CRCA were merged into CMS2 of the consensus classification ^13^.

Recently, the exploratory clinical applicability of our CRCA subtypes was demonstrated when secondary analysis of a randomised clinical trial assessing patient benefit from the addition of oxaliplatin to fluorouracil-leucovorin in early-stage disease revealed that benefits were highly enriched in the enterocyte subtype compared to the other subtypes in the discovery cohort. Therefore, in this study we have applied our CRCA subtype classification for assay development and evaluation.

Translating these findings into routine clinical practice remains challenging, mainly due to the lack of a fit-for-purpose assay that can classify patient samples into subtypes within a clinically relevant turnaround time and reasonable costs using formalin-fixed paraffin-embedded (FFPE) samples. All classifiers were developed from microarray or RNA-seq gene expression profiles, which are expensive, time consuming, require dedicated bioinformatics expertise, and have turnaround times incompatible with clinical applicability. They also rely on pre-amplification of RNA, with consequent impact on accuracy and reproducibility. We previously demonstrated proof-of-concept assays using immunohistochemistry and quantitative reverse transcriptase polymerase chain reaction (qRT-PCR) methods ^3^. Nevertheless, these methods may suffer from reproducibility issues. Hence, we applied nCounter platform (NanoString Technologies) to develop a clinically-relevant biomarker assay for CRC subtype classification.

The nCounter platform has previously been exploited to develop the Food and Drug Administration (FDA)-approved Prosigna^®^ Breast Cancer Prognostic Gene Signature Assay ^15^ to predict risk of recurrence in patients treated with adjuvant hormonal therapy, as well as assays to predict medulloblastoma ^16^ and lymphoma ^17^ subtypes. This platform measures gene expression in the form of discrete counts of barcoded mRNAs, and requires no amplification step, eliminating a potential source of bias. In the present study, we evaluated the suitability of this platform for a gene expression-based assay for our CRCA (a potential surrogate for CMS) subtypes using a modified protocol to subtype CRCs in 4 different cohorts (fresh-frozen and FFPE samples) with different clinical and mutational characteristics. The results were compared to the CMS subtype classification and other platforms. A summary of the classifiers utilised in this study is given in Table 1.

**Table 1.** Summary of the classifiers utilised in this study. Overview of expression profile classifiers: derivation, platform, publication in which it was first introduced, and relationships of the classifiers to each other.

| Classifier | Description | Platform | Original publication |
| --- | --- | --- | --- |
| *786-gene signature* | • 786 genes defining the 5 CRCAssigner (CRCA) subtypes (enterocyte, goblet-like, inflammatory, TA and stem-like)<br>• Derived from gene expression profiles on microarray platform of 387 primary CRC samples.<br>• Potential subtype-specific drug responses have been suggested for stem-like (FOLFIRI) and a subset of TA tumours (cetuximab) using a small cohort of samples (rigorous validation required) <sup>3</sup>, and suggested in the enterocyte subtype (oxaliplatin) by others <sup>14</sup>. | Microarray/<br>RNA-Seq | Sadanandam *et. al.* 2013 <sup>3</sup> |
| *CMS* | • 693 genes defining 4 CMS subtypes (CMS1-4), here applied to microarray/RNA-Seq platforms.<br>• Derived from reconciling the CRCA subtypes with 5 additional sets of gene expression subtypes <sup>8-12</sup>. | Microarray/<br>RNA-Seq | Guinney *et. al.* 2015 <sup>13</sup> |
| *38-gene panel* | • A subset of the 786-gene signature genes. 38 genes able to classify samples into the CRCA subtypes with minimal misclassification errors on microarray platform.<br>• 38 genes were part of the 47 genes selected from 786-gene signature (including all 38-gene panel) plus 3 additional genes not used for subtyping.<br>• Derived from a subset of the samples used to train the 786-gene signature-based classifier that were highly typical of their subtype (see *Methods*). | Microarray/<br>RNA-Seq | Introduced here |
| *NanoCRCA* | • A custom nCounter assay (NanoString Technologies) for 38-gene panel above. Gene expression is profiled using a highly reproducible and modified nCounter assay named as *NanoCRCA*. | nCounter Custom<br>Gene Expression<br>Assay (NanoString<br>Technologies) | Introduced here |

## Results and Discussion

### Analytical development and assessment of CRC subtyping using a reproducible assay

In order to evaluate the applicability of the CRC subtype classification in the clinic, we initially developed a custom nCounter assay using a 50-gene panel, including 47 genes selected from the 786-gene signature and 3 additional genes ^3^ (Table S1; see *Methods*). Initially, we applied a standard protocol (in which biotin labels and molecular barcodes are directly attached to the mRNA probes; Figure 1a) from the manufacturer and tested the performance of the custom nCounter assay using primary fresh frozen tumour RNA obtained from 22 CRCs from two different cohorts (Montpellier and OriGene; Table S2a). The distribution of samples across principal subspace using principal component analysis (PCA; Figure S1a) showed no batch effect between the two cohorts. In addition, hierarchical clustering analysis using nCounter profiles clustered these 22 samples into different groups that potentially represent different subtypes (Figure 1b).

**Figure 1.**
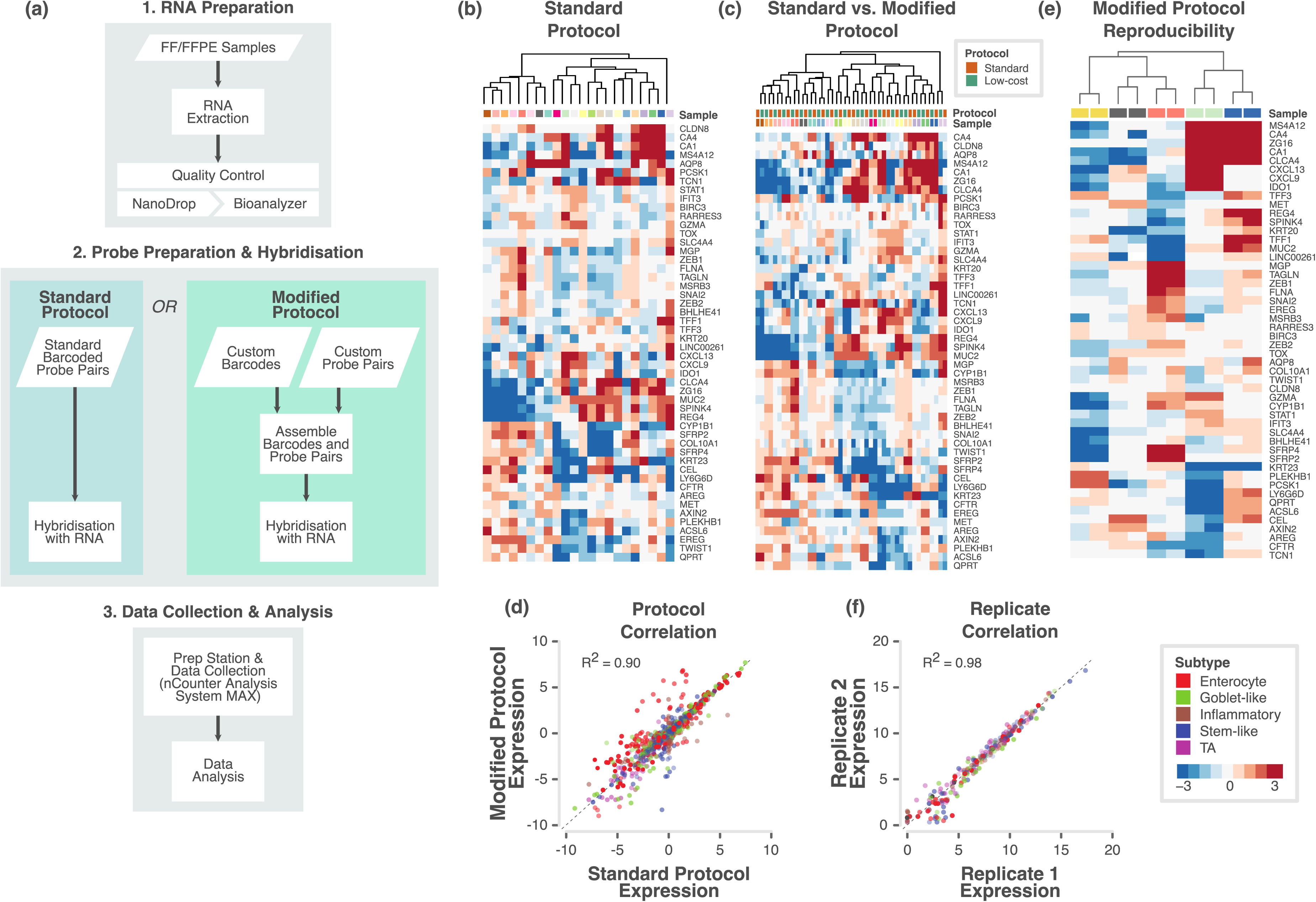
Assessment of different protocols and replicates of reduced subtype gene-based nCounter assay. **a.** Flowchart showing the major steps of the NanoCRCA assay protocols. Specifically, this flowchart demonstrates the difference between standard and modified protocols. Though modified protocol has additional steps, it substantially reduces the cost without significantly increasing the time of the assay. **b-c.** Heatmap of expression levels of the selected 47 subtype-specific genes (and 3 additional genes) for 22 samples from the OriGene and Montpellier cohorts as measured on a custom nCounter panel using b) standard protocol and c) both standard and modified protocols. **d.** A scatter plot of gene expression measurements for all genes in all samples between the standard and modified protocols. Each point is coloured by the gene’s weight (PAM score) in the 786-gene centroids. Correlation co-efficient (R^2^) value is shown. **e.** Heatmap of expression levels of selected 47 subtype-specific genes (and 3 additional genes) from 5 technical replicate samples assayed using modified protocol with a maximum interval of 40 weeks. **f.** Scatter plot of gene expression measurements for all genes in all samples between technical replicates (median centered within data sets before correlation to remove batch effects). Each point is coloured by the gene’s weight (PAM score) in the 786-gene centroids. Correlation co-efficient (R^2^) value is shown.

Next, we evaluated if a “modified” protocol (custom unique probes are attached to biotin labels and molecular barcodes separately; approximately ~35% less expensive than standard protocol; Figure 1a) from NanoString Technologies can deliver similar classification performance compared to the standard protocol-based assay. The results from the modified protocol-based profiling (Table S2b) showed similar distribution of samples (n=22) across principal subspace using PCA (Figure S1b) and clustered into potential subtypes (Figure S1c) in a similar fashion to the standard protocol.

We merged both the standard and modified protocols’ gene expression data after normalising (gene-wise median centring) each dataset and performed PCA (Figure S1d). Figure 1c shows the clustering of the same samples between protocols. Measurements showed high correlation (R^2^=0.90, p<0.001; Figure 1d) between these different protocols. This demonstrates that we can successfully replicate results from standard protocol using modified protocol for a more cost-effective assay. Therefore, we adopted the modified protocol for our assay (Figure 1a).

To test if our assay results are highly reproducible between batches, we performed our assay on five of the above samples twice, in separate batches of maximum 40 weeks apart; Table S2c-d). Figure 1e and Figure S1e shows the clustering of replicate samples together with negligible batch effect. We achieved high concordance between the assays across the two replicates, with a Pearson correlation R^2^ of 0.98 (p<0.001; Figure 1f). This establishes the high reproducibility of our assay over non-negligible periods of time. Hence, in the future, we can use this assay to test the state of subtypes using matched pre- and post-treatment biopsies or surgical materials.

### Analytical validation of CRC subtype assay using FFPE samples

In the clinic, FFPE tumour samples are more prevalent than those that are fresh frozen. Hence, clinical diagnosis is mainly dependent on FFPE samples. At the same time, FFPE samples can present challenges of low abundance or highly degraded RNA. We therefore tested NanoCRCA assay using FFPE samples. Using the FFPE-preserved cohort of samples (see *Methods*), we assessed the congruence of the standard and modified assay protocols (Figures S2a-e and Table S3a-c) and the reproducibility of the modified protocol (Figure S2f and Table S3d-e) in FFPE samples, as previously with fresh frozen samples. We again achieved successful clustering of samples by sample rather than by protocol (Figure 2a). Pearson’s correlation coefficient of gene expression between the standard and modified protocols for 12 samples was 0.88 (Figure 2b). Five pairs of technical replicates also showed highly reproducible results (Figure 2c) with a Pearson’s correlation coefficient of 0.96 (Figure 2d), similar to that for fresh frozen samples (Figure 1f).

**Figure 2.**
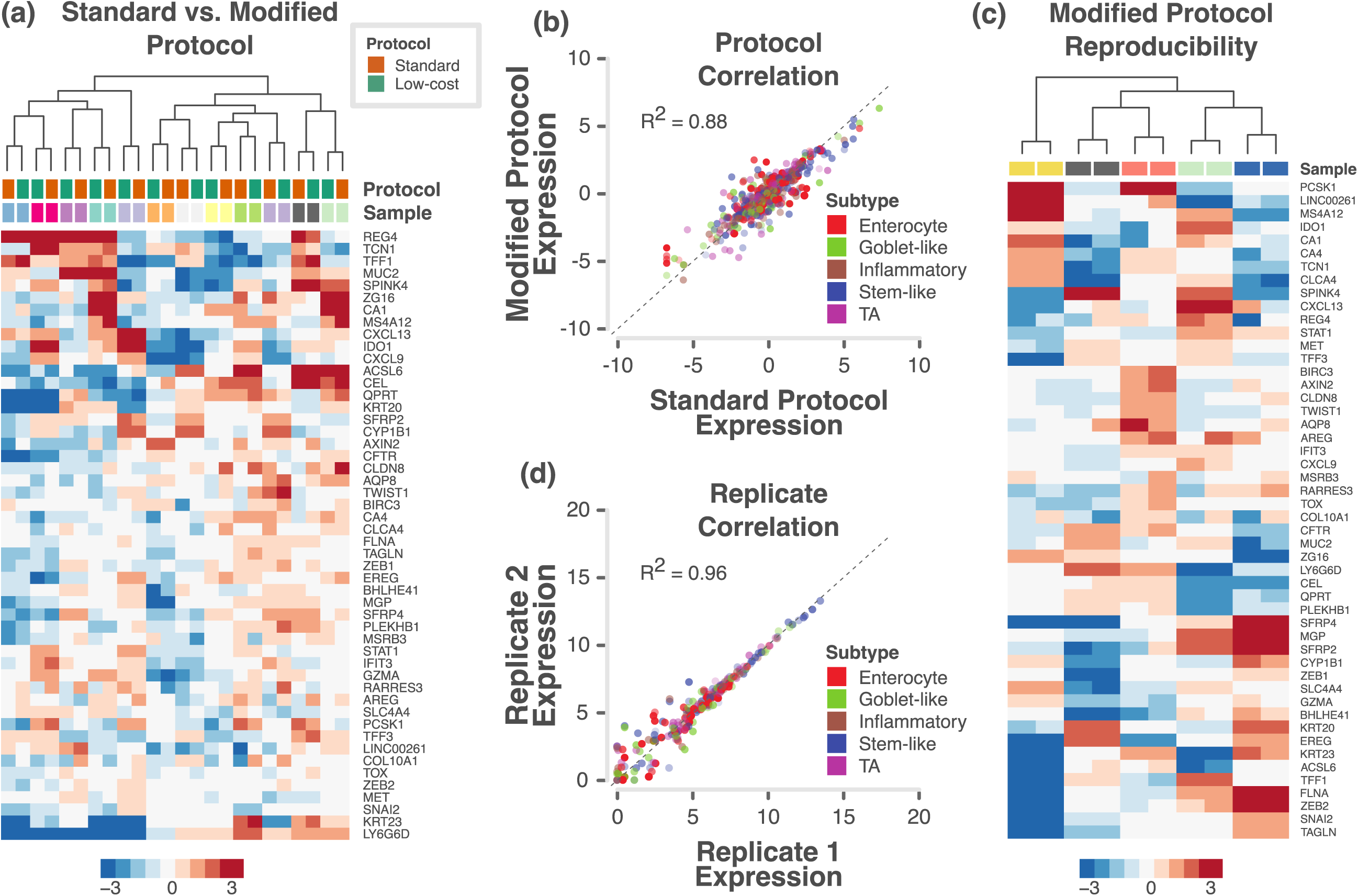
Assessment of protocols and reproducibility using nCounter in FFPE samples. **a.** Heatmap of expression levels of the selected 47 subtype-specific genes (and 3 additional genes) for 24 samples from the RETRO-C cohort as measured on a custom nCounter panel using both standard and modified protocols. **b.** A scatter plot of gene expression measurements for all genes in all samples between the standard and modified protocols. Each point is coloured by the gene’s weight (PAM score) in the 786-gene centroids. Correlation co-efficient (R^2^) value is shown. **c.** Heatmap of expression levels of selected 47 subtype-specific genes (and 3 additional genes) from 5 technical replicate samples assayed using modified protocol with a maximum interval of 13 weeks. **d.** Scatter plot of gene expression measurements for all genes in all samples between technical replicates (median centered within data sets before correlation to remove batch effects). Each point is coloured by the gene’s weight (PAM score) in the 786-gene centroids. Correlation co-efficient (R^2^) value is shown.

### Selection of robust gene set for subtyping

Successful clinical biomarker assays should be able to classify samples into subtypes with high concordance, and this requires a robust set of genes. Hence, we tested the robustness of our selected 47 genes (out of 50 genes from 786-gene signature) using two in-laboratory developed bioinformatics tools – *idSample* and *intPredict* (Figure 3a). Since “mixed subtype” samples with more than one subtype were present in CRC ^13^, we selected only the samples from our published training dataset (n=387) ^3^ that showed at least 70% probability for a single subtype (n=192, Figure 3b; Table S4a). This was done using *idSample* tool that employs support vector machine regression method (*see Methods*). Furthermore, *intPredict* tool, which contains a pipeline of supervised class prediction methods, was used to identify 38 robust genes (38-gene panel) out of 47 genes with lowest percentage misclassification error rate (MCR, 1%; Figure 3c-d; Table S4b-c; *Methods*). In order to further effectively classify our samples into CRC subtypes using the selected 38-gene panel, we calculated newly-derived 38-gene panel classification scores (gene centroids using Prediction Analysis of Microarrays methodology; see *Methods*; Figure 3e) having only 1.6% MCR to classify samples profiled on our nCounter assay, and compared to the other gene profiling platforms and classifiers in three different CRC cohorts.

**Figure 3.**
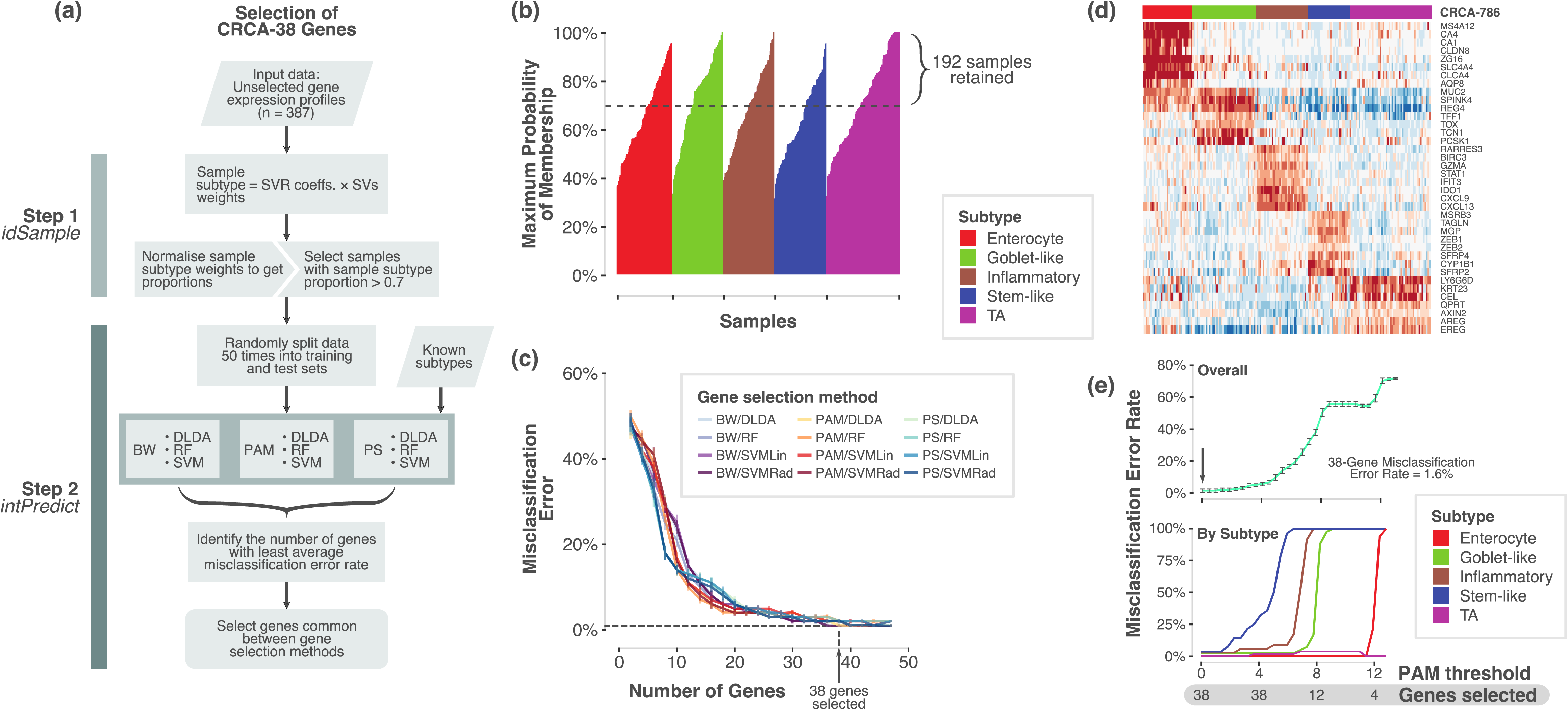
Selection of robust 38-gene panel. **a.** Overview of the process and pipelines used to select a robust gene set for the NanoCRCA assay. **b.** A bar plot showing samples (n=387) from our original published dataset ^3^ that shows probability of a sample belonging to a given subtype as assessed using *idSample* tool developed by us. **c**. A line plot showing MCR and number of genes as modelled using intPredict pipeline of class prediction methods and samples from b). **d.** Heatmap showing the gene expression of the 38-gene panel selected by the pipeline in the 192 selected samples from b). Top bar shows the 786-gene signature-based subtype of the samples. **e**. Line plots showing MCR using PAM analysis at different gene selection and PAM thresholds for all the subtypes (upper) and individual subtypes (lower).

### Subtyping of fresh frozen samples

To determine if these assays could successfully stratify patient samples into CRCA subtypes, we applied our NanoCRCA assay (utilising the modified protocol and 38-gene signature) to fresh frozen CRC samples (n=57; combined samples from the Montpellier, Singapore and OriGene cohorts). Subtypes were determined by the correlation of gene expression profiles with the 38-gene panel centroids. All five subtypes were identified by the assay and demonstrated distinct patterns of gene expression (Figure 4a). A proportion of samples (9/57) were determined to be of mixed subtype (Figure 4a; Table S5), indicating the presence of multiple distinct subtypes within the same tumour, as previously demonstrated^13^.

**Figure 4.**
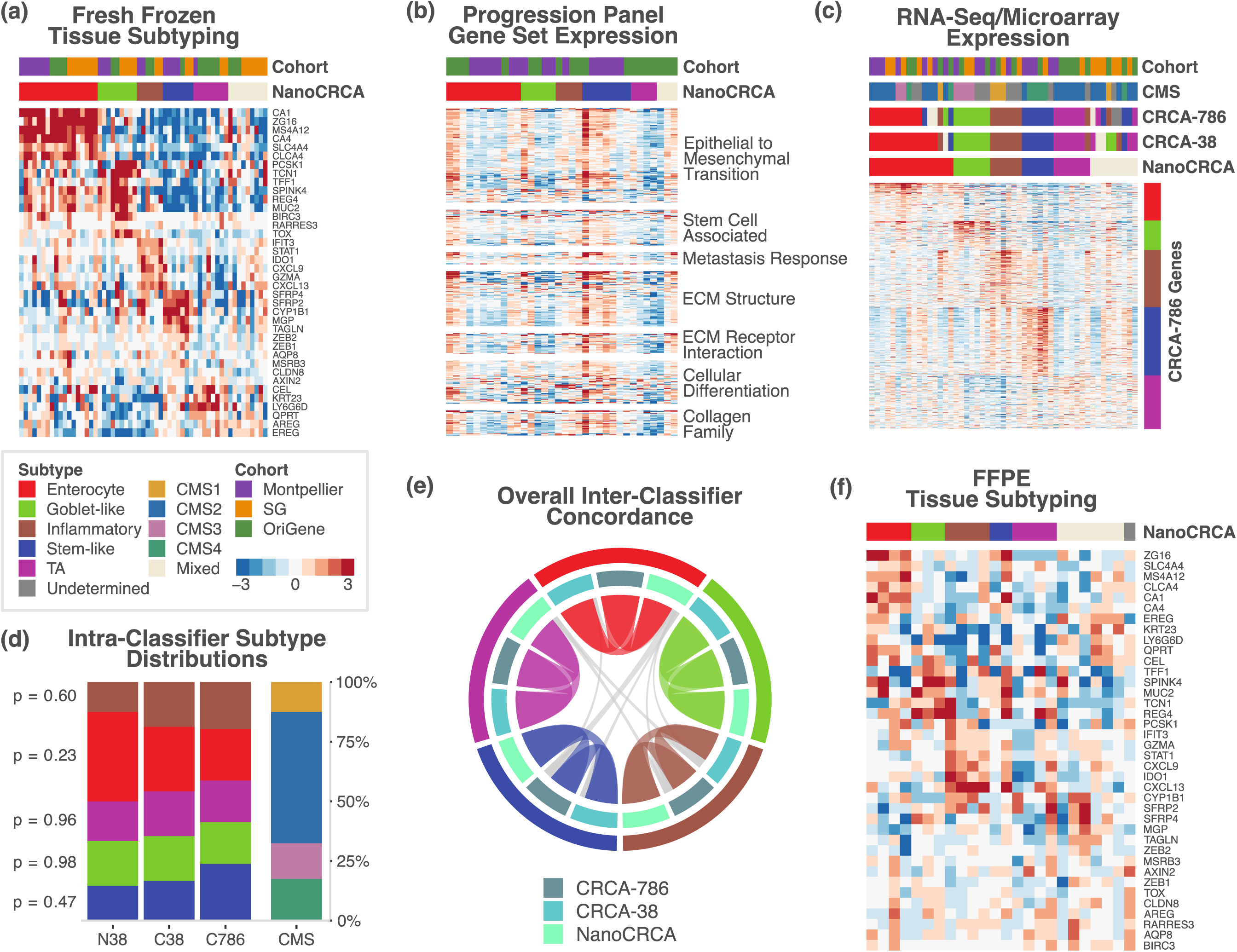
NanoCRCA subtyping, pathway and 786-gene signature analysis with subtype stability. **a.** Heatmap showing the expression of 38-gene panel in the three fresh frozen cohorts as measured using NanoCRCA assay. The lower top bar indicates the NanoCRCA subtype of the samples and the upper top bar indicates the cohorts. **b.** Heatmap of nCounter PanCancer Progression Panel-based gene expression profiles from the Montpellier and OriGene cohorts of samples (n=34). The lower top bar indicates the NanoCRCA subtype of the samples and the upper top bar indicates the cohorts. Genes are grouped according to functional annotations provided by NanoString Technologies or subtype gene signatures. **c.** Heatmap of batch-corrected RNA-Seq/microarray gene expression profiles from all the three cohorts (n=51). The NanoCRCA subtypes, 38-gene panel, 786-gene signature, CMS and different cohorts of the samples are shown from bottom to the top bars respectively. Subtype-specific gene signatures representing each subtype are shown on the side. **d.** Distribution of subtypes according to each classifier. Samples that were mixed or undetermined subtype were excluded for each classifier. Results of statistical tests of proportion between the three subtype-specific classifiers are shown on the left-hand side. **e.** Chord plot illustrating the tendency of samples to be classified as the same subtype between the three subtype-based assays. Samples from all three cohorts which had no mixed or undetermined subtype calls were included (n=39). Each arc connects the classification of a sample in two different assays, and each sample is represented by three arcs (connecting NanoCRCA assay (nCounter platform), 38-gene panel and 786-gene signature (microarray platform) subtypes). Samples with the same subtype in all three assays are coloured by their subtype. Samples (only four of 39 samples) that had discordant classification between the assays are coloured grey. **f.** Heatmap showing the expression of 38-gene panel genes in FFPE samples as measured using NanoCRCA assay. The subtypes as assigned using NanoCRCA assay are shown on the top bar.

### Biological characteristics of the identified subtypes

To further confirm the molecular characteristics of the subtypes, we performed analysis using NanoString Technologies’ PanCancer Progression Panel (Table S6a-b; Figure S3a-b). As expected, the stem-like samples had increased expression of genes associated with epithelial to mesenchymal transition (EMT), stem cells, metastatic response, extracellular matrix (ECM) structure and receptor interaction, cellular differentiation, collagen family and others (Figures 4b and S3c-d). In addition, the expression of the 786-gene signature is shown in Figure 4c alongside the NanoCRCA subtypes of the samples. Overall, these analyses demonstrate that the subtypes identified by NanoCRCA assay represent the published^3^ molecular characteristics of these subtypes in 51 samples (with matched gene expression profiles from other platforms) from three independent cohorts.

### Concordance and distribution of subtypes across fresh frozen samples between platforms

We sought to confirm that subtyping using the NanoCRCA assay (with the modified protocol and 38-gene signature) mirrored the results of subtyping using highly multiplexed platforms such as microarrays and RNA-Seq. Matched microarray or RNA-Seq data for the fresh-frozen Montpellier, Singapore and OriGene cohorts were generated or downloaded from public repositories (see *Methods*) (n=51). Subtypes were determined by correlation of the gene expression profiles with both our original 786-gene signature centroids^3^ and our new 38-gene panel centroids.

As a confirmation that platform and gene set differences did not bias the distribution of subtypes assigned to the samples, Figure 4d shows there is no significant difference (p>0.05; proportion tests) in the distribution of each subtype across the two CRCA classifiers from different platform and CMS classifier in the three cohorts. This again validates the similarity between CRCA and CMS subtypes.

To assess the stability of individual subtypes across the various platforms and gene sets, we plotted Figure 4e using subtype-determined and non-mixed samples for these three fresh frozen cohorts (n=39; Table S5). The three assays are shown (786-gene signature, 38-gene panel and NanoCRCA) along with the 5 CRCA subtypes. While the goblet-like subtype was consistent across all the different classifications, there were only four samples that had different subtypes across classifications (shown in grey in Figure 4e). Two of the samples were classified as either enterocyte or stem-like, one as either enterocyte or inflammatory, and one as either TA or inflammatory. Three of these four samples were from the OriGene cohort, potentially due to platform-specific effects as discussed below. However, overall concordance between platforms was good (Figure 4e), with 35 of 39 non-mixed/undetermined samples (89.7%) showing the same subtype across all 3 assays.

### Subtyping of FFPE-preserved samples

Applying the NanoCRCA assay to FFPE samples from our own clinical cohort (see *Methods*; n=24) revealed that all the CRCA subtypes were present in this cohort along with mixed and undertermined samples (Figure 4f and Table S3e-f). Regardless of the generally lower quality of RNA obtained from FFPE samples, this pilot cohort demonstrates that the NanoCRCA assay can be used to determine the subtypes of FFPE-preserved samples, thereby making our assay widely practicable in a clinical setting.

Assessment of CRC subtypes in Montpellier cohort using NanoCRCA assay

In order to understand the clinical significance of stratifying CRC samples using our NanoCRCA, we further analysed our Montpellier cohort of 17 primary tumours (stage IV) patients ^4, 6^ (*Methods*; Figures 5a and S4a-b; Table S7a-b). All the CRCA subtypes were present in this cohort, and all samples were successfully classified, with none showing mixed or undetermined subtype characteristics (Figure 5b-c). We observed a non-uniform distribution of the subtypes with enterocyte contributing 41.2% of all the samples followed by stem-like (23.5%) and goblet-like (17.6%) subtypes. However, inflammatory (11.8%) and TA (5.9%) subtypes were low in numbers in this cohort of samples (Figure 5c).

**Figure 5.**
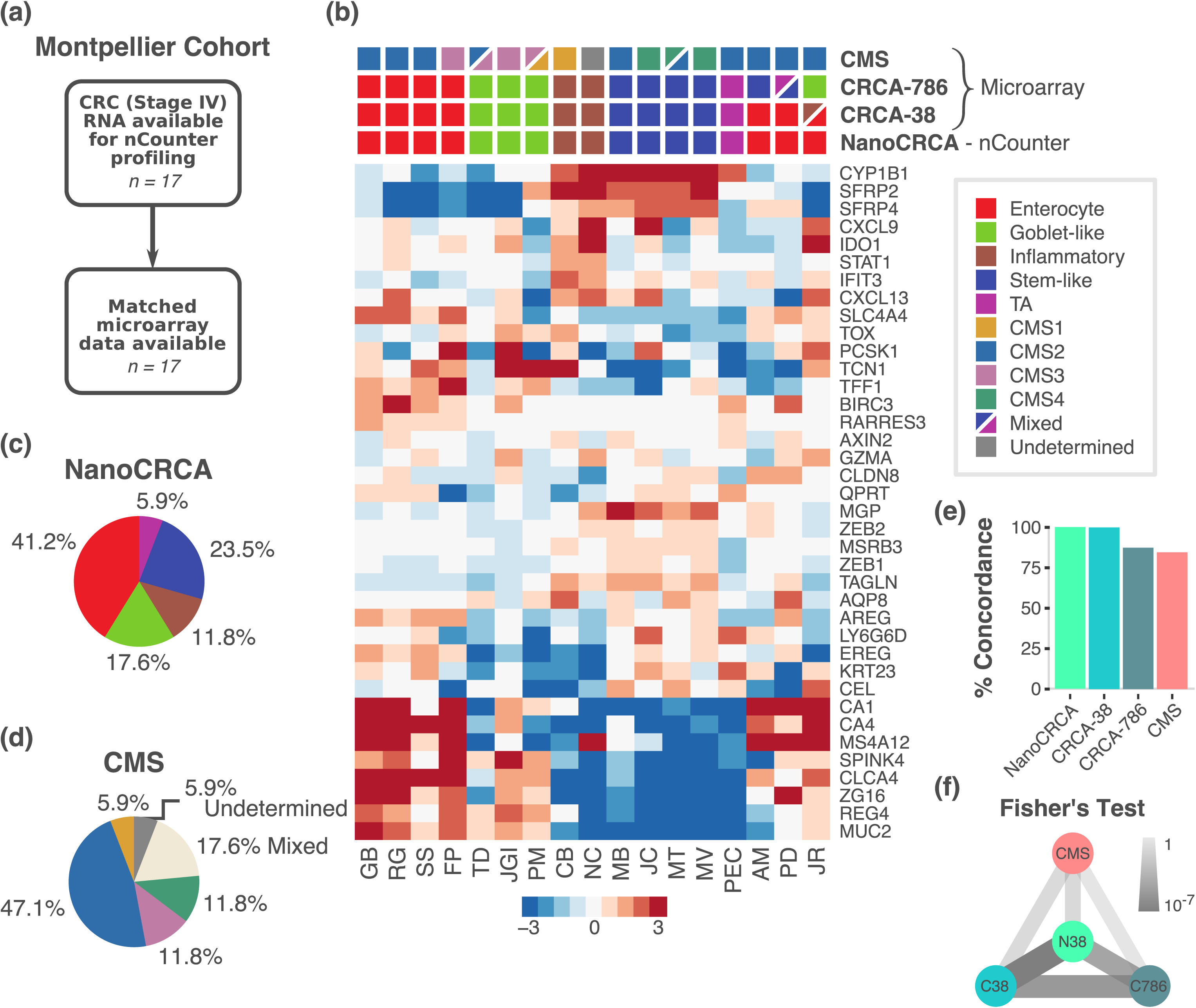
Montpellier cohort - NanoCRCA assay, its comparison with other platforms and CMS classifier and characteristics. **a.** A summary of the Montpellier cohort showing patient characteristics, sample size and available HG-U133 Plus 2.0 microarray data. **b.** Heatmap showing the expression of 38-gene panel in the Montpellier cohort as measured using NanoCRCA assay. The subtypes as assigned using NanoCRCA assay (nCounter platform), 38-gene panel, 786-gene signature and CMS classifications (microarray platform) are shown on the top bars. **c**. Pie chart showing the proportion of different subtypes (including mixed and undetermined samples) from NanoCRCA classification. **d**. Pie chart showing the proportion of different subtypes (including mixed and undetermined samples) from CMS classification. **e-f.** Comparisons between NanoCRCA and other classifications including 38-gene panel, 786-gene signature and CMS showing e) percent concordance and f) statistical significance (Fisher’s exact test).

### Comparison of subtypes with CMS classifier and across platforms

We additionally compared our NanoCRCA assay with the microarray-based CMS subtypes in this cohort. We classified the Montpellier cohort of 17 microarray gene expression profiles into CMS subtypes using the published CMS classifier^13^. We successfully classified the samples into all of the CMS subtypes: 47.1% were CMS2 (enterocyte and TA); 11.8% each of CMS3 (goblet-like) and CMS4 (stem-like); and 5.9% of CMS1 (inflammatory). However, we found 17.6% mixed and 5.9% undetermined samples (Figure 5d). Thus, NanoCRCA showed good concordance (84.6%) with the CMS classifier excluding the mixed and undetermined samples (Figure 5e). We further performed pairwise Fisher’s exact test between the CMS and NanoCRCA subtypes (Figure 5f), which confirmed borderline significant association to the CMS classification (FDR=0.07). This suggests that NanoCRCA may be applied as a surrogate to predict CMS subtypes (as CMS classification was partly derived from CRCA classification) with reasonable concordance in addition to CRCA subtypes.

We also compared the performance of the NanoCRCA assay against classification using Affymetrix Human Genome U133 Plus 2.0 (HG-U133 Plus2) microarray profiles ^4^ in this cohort and the new 38-gene signature (Figures 5b and S4c). We classified the samples’ microarray profiles into all the five subtypes with 35.2% as enterocyte, 23.5% as stem-like, 17.6% as goblet-like, 11.8% as inflammatory and 5.9% as TA (Figure S4d). Only one sample was defined as a mixed subtype (1/17, 5.9%) expressing both inflammatory and enterocyte genes. The expression of enterocyte genes was consistent with the classification of the sample as enterocyte on the NanoCRCA platform. The fact that the sample was classified as a mixture of subtypes by the 38-gene panel may be attributed to platform-specific effects. Overall, the NanoCRCA assay showed perfect 100% concordance with the microarray-based 38-gene panel classification after excluding samples with mixed classification (due to challenges in comparing these mixed subtypes to others) (Figure 5e).

Similarly, we compared the 786-gene signature-based classification using microarray data to the results of NanoCRCA. The 786-gene signature-based classification yielded 23.5% enterocyte, 29.4% stem-like, 23.5% goblet-like, 11.8% inflammatory and 5.9% TA samples (Figure S4e). There was one mixed sample (1/17, 5.9%; Figures 5b and S4e) that was different from that called mixed by 38-gene panel. Irrespective of the different number of genes profiled on the different platforms, the NanoCRCA assay showed 87.5% concordance with the 786-gene signature subtypes (Figure 5e). Again, the 12.5% discordance may be attributable to noisy genes present in the 786-gene signature. Overall, the NanoCRCA assay and 786-gene signature classification perform well with good concordance.

We statistically validated these findings by applying Fisher’s exact test on these classifications, excluding mixed or undetermined samples. We found that the NanoCRCA assay was significantly (false discovery rate; FDR<0.001; Figure 5f) associated with both the 38-gene panel and 786-gene signature classification systems. This again statistically validates the high concordance between NanoCRCA classification and different gene- and platform-based CRCA classifications, further confirming the robustness of our NanoCRCA subtypes. In sum, all the three classifications from the two different platforms identified all the five subtypes and the NanoCRCA assay predicted robust subtypes highly consistent with the microarray platform.

### NanoCRCA assay-based CRC subtypes and associated mutational profiles in a multi-stage Asian cohort

Our profiling of the multi-stage SG cohort using NanoCRCA is the first of its kind in an Asian population, to our knowledge. We further observed high concordance in subtypes between the NanoCRCA assay (n=23) and RNAseq (for both 38-gene panel and 786-gene signature; n=17; Figures S5a-e; Table S7c-d). There was no visible trend in association between stages and subtypes (Figure S5f; Table S7e).

Previously, we reported that the inflammatory (CMS1) subtype is associated with MSI and *BRAF* mutations, whereas goblet-like (CMS3) subtype is highly associated with *KRAS* mutations ^3, 13, 18^. To further validate the NanoCRCA subtypes, we compared these with the mutational (*BRAF* and *KRAS*) and MSI status (Figure 6a and Table S7e) of cancers in the Asian cohort (n=11) that were detected within our laboratory using methods described in *Supplementary Information*. All (100%) the inflammatory subtype CRCs were associated with MSI. Interestingly, one of the two *BRAF* mutant tumours was associated with the inflammatory (CMS1) subtype and MSI status. Similarly, all three goblet-like subtype (CMS3; 100%) tumours were associated with *KRAS* mutation. There were three other *KRAS* mutant tumours associated with the enterocyte or stem-like subtypes, representing *KRAS* mutant tumours are also associated less frequently with other subtypes, as previously reported^13^. This analysis with our NanoCRCA assay corresponds with known associations of subtypes with mutational and MSI profiles with this small SG data set of 17 samples. However, additional large data sets and NanoCRCA assays are warranted to validate these observations.

**Figure 6.**
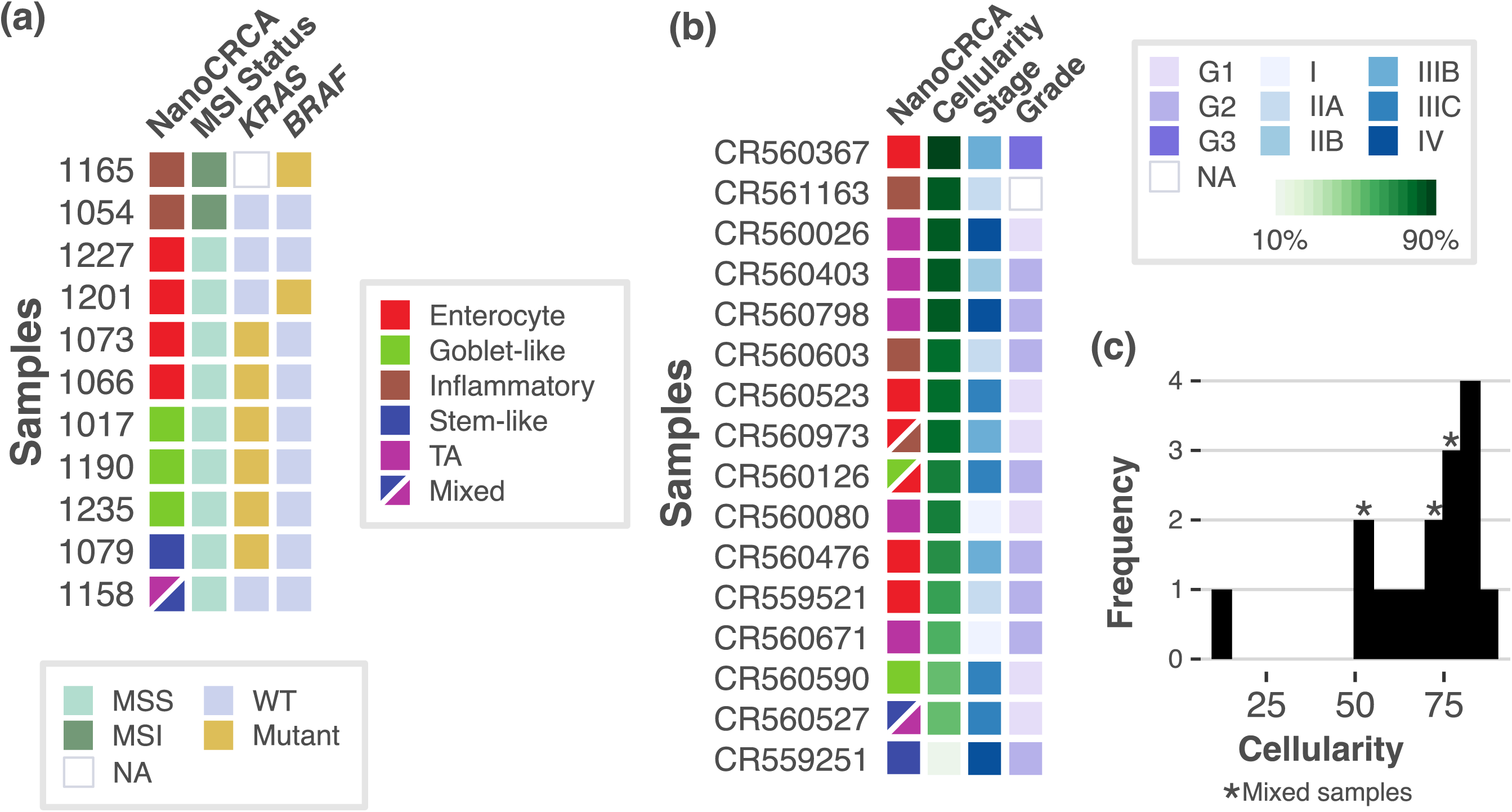
Subtype association with MSI and mutations and effect of tumour cellularity. **a.** A plot showing MSI and mutational (*KRAS* and *BRAF*) status of samples alongside subtypes from NanoCRCA classification using SG cohort. **b.** A plot showing tumour cellularity, stage and grade associated with subtypes from this classification using OriGene cohort. **c.** Histogram showing the distribution of tumour cellularity using OriGene cohort.

### Effect of tumour cellularity on CRC subtyping using NanoCRCA assay

Next, we sought to test if tumour cellularity affects our NanoCRCA assay using OriGene cohort of samples (n=17) spanning all stages of CRC (Figures 6b and S6a-d; Table S7f-g). We observed high concordance between NanoCRCA assay and HTA microarray (for both 38-gene panel and 786-gene signature; Figure S6e), although CA1 gene was not present in the HTA array and there was a low correlation (R^2^<0.5) in expression of two inflammatory genes between HTA and NanoCRCA assay (Figure S6f and Supplementary Information).

With variable tumour cellularity ranging from 10% to 85% in this cohort, we postulated that if cellularity affects our assay, the low cellularity samples should be either qualified as “undetermined” or “mixed subtype” samples. One sample had extremely low cellularity of 10% and one additional sample was from a liver metastasis. Interestingly, none of the samples were classified as having undetermined subtype, regardless of cellularity. On the other hand, the three mixed samples had varying levels of cellularity (50-75%) (Figures 6b-c). These results suggest that our NanoCRCA may not be affected by cellularity due to the selection of robust gene sets, which requires further validation in the future.

## Conclusion

In summary, we analytically developed and validated NanoCRCA biomarker assay based on robust 38-gene panel and classified CRC samples into molecular subtypes, along with mixed subtypes. Subtype prediction by the NanoCRCA assay is highly concordant with CMS subtypes and multiple platforms. Also, stratification using this assay reproduces the clinical, mutational and molecular characteristics of the CRC subtypes. Since multiple CRC clinical trials require low-cost, reproducible and rapid clinically implementable assays to prospectively validate CRC subtypes for subtype-specific studies, our NanoCRCA assay may potentially facilitate this process in the clinic using FFPE samples.

## Methods

### Patient cohorts

Four CRC cohorts were studied: three derived from fresh-frozen and one from FFPE samples. The first included 17 metastatic (stage IV) CRCs (Montpellier cohort) from patients with no prior chemotherapy from a published study ^4^ (microarray data at GEO accession GSE62080). A second cohort (Singapore; SG) included 23 untreated CRC samples from patients (20 Chinese, 2 Malay and 1 Indian) participating in an on-going observational CRC study from the National Cancer Centre of Singapore and Singapore General Hospital (SingHealth Institutional Review Board IRB: 2013/110/B). RNA-Seq for these patients were performed (*Supplementary Methods*) and data are deposited with GEO accession GSE101588. A third cohort (OriGene; n=17) was purchased from OriGene (Rockville, MD, USA) (microarray data at GEO accession GSE101472). The final cohort consisted of 24 FFPE CRC samples from “A retrospective translational study at The Royal Marsden NHS Foundation Trust: characterisation of molecular predictors of response to cetuximab or panitumumab in patients with colorectal cancer (RETRO-C)”, with IRB and ethical approval (NRES Committee East of England-Cambridge Central, 10/H0308/28).

### Subtype concordance and significance

Subtype concordance between two different platforms was calculated as the percentage of samples that showed the same subtype (not including mixed and undetermined samples) in both. Subtypes were deemed concordant between CRCA and CMS subtypes based on the following equivalence: CMS1=Inflammatory; CMS2=Enterocyte and TA; CMS3=Goblet-like; CMS4=Stem-like ^13^. Fisher’s exact test between different platforms or classifiers was performed to assess statistical significance of their subtype concordance, and p-values adjusted using the FDR method.

*Supplementary Methods & Materials* contains detailed information of all methods employed.

## DATA ACCESS

Previously published GEO Omnibus data sets were analysed for gene set selection (GSE14333 and GSE13294) and microarray-based subtyping of the Montpellier cohort (GSE62080). nCounter data for all cohorts (GSE101479 – standard protocol and GSE101481 – modified protocol) and microarray/RNA-Seq data for OriGene (GSE101472) and Singapore (GSE101588) cohorts are deposited under the SuperSeries with accession number GSE101651.

## ETHICAL APPROVAL AND INFORMED CONSENT

The protocol for the SG cohort of this study was approved by the SingHealth Institutional Review Board: 2013/110/B. The protocol for the RETRO-C cohort was approved by NRES Committee East of England-Cambridge Central: 10/H0308/28. Montpellier cohort had ethical approval as described in their original publication and OriGene cohort is available from the commercial vendor.

## ACKNOWLEDGEMENTS

We thank Prof. Mitch Dowsett, Dr. Richard Buus, Dr. Maggie Cheang, Dr. Nicola Valeri, Dr. George Vlachogiannis and Dr. Andrea Lampis from the ICR for their advice on the use of nCounter platform, and Prof. Paul Workman for helpful comments on the manuscript. We thank Rosetrees Trust’s grant for consumables (Reference Number: A1255). We acknowledge NHS funding to the NIHR Biomedical Research Centre at The Royal Marsden and the ICR.

## AUTHOR CONTRIBUTIONS

A.Sa. conceived the idea, designed the experimental/bioinformatics strategies, nCounter assay protocols and gene sets, performed a part of the bioinformatics analysis, and supervised the entire project. C.R. optimised, developed and performed nCounter assays, and performed microarray experiments. K.E. performed bioinformatics analysis of all types of data and prepared the figures. C.R and E.F. performed RNA extraction and modified protocol NanoCRCA assays on RETRO-C cohort FFPE samples. G.N. developed class prediction models and the *R* package. Y.P. performed RNA-Seq data processing and data integration. P.P. assisted with data analysis. M.D.R., and P.M. isolated and provided RNA for Montpellier cohort of samples and their associated clinical information and microarray data; and K.S., T.W.S., and I.B.T., isolated and provided RNA for SG cohort and their associated RNA-Seq data and clinical information. R.T.L., and A.Sc. helped with manuscript preparation. L.S.T.M. prepared FFPE slides for RNA extraction in the RETRO-C cohort; F.S., and D.C., designed the RETRO-C study protocol and R.B. collected and prepared the tissue samples. C.R., K.E., and A.Sa. interpreted the results and wrote the manuscript.

## DISCLOSURE DECLARATION

A.Sa. has ownership interest (including patents) as a patent inventor for a patent entitled “Colorectal cancer classification with differential prognosis and personalized therapeutic responses” (patent number PCT/IB2013/060416).

## References

1. Ferlay J, Soerjomataram I, Ervik M, Dikshit R, Eser S, Mathers C, Rebelo M, Parkin D, Forman D, Bray F. Cancer incidence and mortality worldwide: sources, methods and major patterns in GLOBOCAN 2012. International Journal of Cancer 2015;136.

2. Van Cutsem E, Cervantes A, Adam R, Sobrero A, Van Krieken JH, Aderka D, Aguilar EA, Bardelli A, Benson A, Bodoky G. ESMO consensus guidelines for the management of patients with metastatic colorectal cancer. Annals of Oncology 2016;27: 1386–422.

3. Sadanandam A, Lyssiotis CA, Homicsko K, Collisson EA, Gibb WJ, Wullschleger S, Ostos LCG, Lannon WA, Grotzinger C, Del Rio M, Lhermitte B, Olshen AB, et al. A colorectal cancer classification system that associates cellular phenotype and responses to therapy. Nature Medicine 2013;19: 619–25.

4. Del Rio M, Molina F, Bascoul-Mollevi C, Copois V, Bibeau F, Chalbos P, Bareil C, Kramar A, Salvetat N, Fraslon C, Conseiller E, Granci V, et al. Gene expression signature in advanced colorectal cancer patients select drugs and response for the use of leucovorin, fluorouracil, and irinotecan. Journal of Clinical Oncology 2007;25: 773–80.

5. Khambata-Ford S, Garrett CR, Meropol NJ, Basik M, Harbison CT, Wu S, Wong TW, Huang X, Takimoto CH, Godwin AK, Tan BR, Krishnamurthi SS, et al. Expression of epiregulin and amphiregulin and K-ras mutation status predict disease control in metastatic colorectal cancer patients treated with cetuximab. Journal of Clinical Oncology 2007;25: 3230–7.

6. Del Rio M, Mollevi C, Bibeau F, Vie N, Selves J, Emile J-F, Roger P, Gongora C, Robert J, Tubiana-Mathieu N, Ychou M, Martineau P. Molecular subtypes of metastatic colorectal cancer are associated with patient response to irinotecan-based therapies. European Journal of Cancer 2017;76: 68–75.

7. Medico E, Russo M, Picco G, Cancelliere C, Valtorta E, Corti G, Buscarino M, Isella C, Lamba S, Martinoglio B, Veronese S, Siena S, et al. The molecular landscape of colorectal cancer cell lines unveils clinically actionable kinase targets. Nature Communications 2015;6: 7002.

8. De Sousa E Melo F, Wang X, Jansen M, Fessler E, Trinh A, de Rooij LPMH, de Jong JH, de Boer OJ, van Leersum R, Bijlsma MF, Rodermond H, van der Heijden M, et al. Poor-prognosis colon cancer is defined by a molecularly distinct subtype and develops from serrated precursor lesions. Nature Medicine 2013;19: 614–8.

9. Marisa L, de Reyniès A, Duval A, Selves J, Gaub MP, Vescovo L, Etienne-Grimaldi MC, Schiappa R, Guenot D, Ayadi M, Kirzin S, Chazal M, et al. Gene expression classification of colon cancer into molecular subtypes: characterization, validation, and prognostic value. PLoS Medicine 2013;10.

10. Budinska E, Popovici V, Tejpar S, D’Ario G, Lapique N, Sikora KO, Di Narzo AF, Yan P, Graeme Hodgson J, Weinrich S, Bosman F, Roth A, et al. Gene expression patterns unveil a new level of molecular heterogeneity in colorectal cancer. Journal of Pathology 2013;231: 63–76.

11. Schlicker A, Beran G, Chresta CM, McWalter G, Pritchard A, Weston S, Runswick S, Davenport S, Heathcote K, Castro DA, Orphanides G, French T, et al. Subtypes of primary colorectal tumors correlate with response to targeted treatment in colorectal cell lines. BMC Medical Genomics 2012;5: 1–15.

12. Roepman P, Schlicker A, Tabernero J, Majewski I, Tian S, Moreno V, Snel MH, Chresta CM, Rosenberg R, Nitsche U, Macarulla T, Capella G, et al. Colorectal cancer intrinsic subtypes predict chemotherapy benefit, deficient mismatch repair and epithelial-to-mesenchymal transition. International Journal of Cancer 2013;134: 552–62.

13. Guinney J, Dienstmann R, Wang X, de Reyniès A, Schlicker A, Soneson C, Marisa L, Roepman P, Nyamundanda G, Angelino P, Bot BM, Morris JS, et al. The consensus molecular subtypes of colorectal cancer. Nature Medicine 2015;21: 1350–6.

14. Song N, Pogue-Geile KL, Gavin PG, Yothers G, Rim Kim S, Johnson NL, Lipchick C, Allegra CJ, Petrelli NJ, O’Connell MJ, Wolmark N, Paik S. Clinical outcome from oxaliplatin treatment in stage II/III colon cancer according to intrinsic subtypes: Secondary analysis of NASBP C-07/NRG oncology randomized clinical trial. JAMA Oncology 2016;2: 1162–9.

15. Wallden B, Storhoff J, Nielsen T, Dowidar N, Schaper C, Ferree S, Liu S, Leung S, Geiss G, Snider J, Vickery T, Davies SR, et al. Development and verification of the PAM50-based Prosigna breast cancer gene signature assay. BMC Medical Genomics 2015;8: 54.

16. Northcott PA, Shih DJH, Remke M, Cho YJ, Kool M, Hawkins C, Eberhart CG, Dubuc A, Guettouche T, Cardentey Y, Bouffet E, Pomeroy SL, et al. Rapid, reliable, and reproducible molecular sub-grouping of clinical medulloblastoma samples. Acta Neuropathologica 2012;123: 615–26.

17. Scott DW, Wright GW, Williams PM, Lih C-J, Walsh W, Jaffe ES, Rosenwald A, Campo E, Chan WC, Connors JM, Smeland EB, Mottok A, et al. Determining cell-of-origin subtypes of diffuse large B-cell lymphoma using gene expression in formalin-fixed paraffin-embedded tissue. Blood 2014;123: 1214–7.

18. Poudel P, Nyamundanda G, Ragulan C, Lawlor R, Das K, Tan P, Scarpa A, Sadanandam A. Revealing unidentified heterogeneity in different epithelial cancers using heterocellular subtype classification. bioRxiv 2017.

19. Tibshirani R, Hastie T, Narasimhan B, Chu G. Diagnosis of multiple cancer types by shrunken centroids of gene expression. Proceedings of the National Academy of Sciences 2002;99: 6567–72.

20. Cortes C, Vapnik V. Support-vector networks. Machine Learning 1995;20: 273–97.

21. Newman AM, Liu CL, Green MR, Gentles AJ, Feng W, Xu Y, Hoang CD, Diehn M, Alizadeh AA. Robust enumeration of cell subsets from tissue expression profiles. Nature Methods 2015;12: 453–7.

22. The Cancer Genome Atlas Network. Integrated Genomic Analyses of Ovarian Carcinoma. Nature 2011;474: 609–15.

23. Golub TR, Slonim DK, Tamayo P, Huard C, Gaasenbeek M, Mesirov JP, Coller H, Loh ML, Downing JR, Caligiuri MA. Molecular classification of cancer: class discovery and class prediction by gene expression monitoring. Science 1999;286: 531–7.

24. Dudoit S, Fridlyand J, Speed TP. Comparison of discrimination methods for the classification of tumors using gene expression data. Journal of the American Statistical Association 2002;97: 77–87.

25. Breiman L. Random forests. Machine Learning 2001;45: 5–32.

26. Meyer D, Dimitriadou E, Hornik K, Weingessel A, Leisch F, Chang C-C, Lin C-C, Meyer MD. Package ‘e1071’, 2017.

27. Liaw A, Wiener M. Classification and regression by randomForest. R news 2002;2: 18–22.

28. Dudoit S, Yang Y, Bolstad B. sma: Statistical microarray analysis. https://cranr-projectorg/package=sma 2011.

29. Nielsen T, Wallden B, Schaper C, Ferree S, Liu S, Gao D, Barry G, Dowidar N, Maysuria M, Storhoff J, Henry N, Hayes D, et al. Analytical validation of the PAM50-based Prosigna Breast Cancer Prognostic Gene Signature Assay and nCounter Analysis System using formalin-fixed paraffin-embedded breast tumor specimens. BMC Cancer 2014;14: 177.

